# A Syntaxin1A protein mutation drives sleep reduction in *Drosophila*

**DOI:** 10.1101/2025.09.29.679067

**Authors:** Giovanni Frighetto, Nicola Cellini, Giulio Maria Menti, Mauro Zordan, Aram Megighian

**Author notes:** current affiliation: Department of Integrative Biology and Physiology, UCLA, Los Angeles, 90095, USA.

## Abstract

Syntaxin1A is a key component in the regulation of the vesicle fusion nanomachine and, together with VAMP/synaptobrevin and SNAP-25, forms the core of the SNARE complex, that is the molecular nanomachine regulating neurotransmitter release. Syntaxin1A is localized on the cytosolic face of the presynaptic membrane and with Munc-18-1 is thought to be the starting point for the SNARE complex assembly.^1^ In flies, aspartic-to-arginine amino acid substitution at position 253 of the protein sequence reduces synaptic release by affecting the assembly of multiple SNARE complexes into a super-complex without interfering with the single complex formation per se.^2^ In this study, we analyzed sleep behavior in flies expressing this mutated isoform of Syntaxin1A panneuronally. According to the synaptic homeostasis hypothesis,^3^ the overall decrease of synaptic activity during the previous period of wakefulness should reduce the need for sleep.^4^ We observed a reduction in the total amount of sleep in flies expressing the mutated isoform compared to controls. Moreover, the sleep pattern of the Syntaxina1A mutant flies was more fragmented compared to controls. Video-tracking analysis of free walking flies in an open-field arena showed that these changes were not caused by higher locomotor activity during the daytime relative to controls. Our results suggest that the widespread effect of Syntaxin1A mutation on synapses may lead to a dampening of the information load among neurons and, consequently, impact the regulation of sleep at the cellular level.

**Highlights:**

- The neurotransmitter release machinery modulates sleep in *Drosophila*
- A Syntaxin1A protein mutation causes fragmented night sleep
- Motor activity is not affected by the Syntaxin1A mutation
- Syntaxin1A could regulate the homeostasis of vesicle fusion and sleep

**In brief:** Frighetto et al., investigated the role of a single amino acid substitution in the Syntaxin1A protein in sleep and locomotor behavior. Following the pan-neuronal expression of this mutated Syntaxin1A, they found a specific reduction of sleep with locomotion remaining largely unaffected.

## RESULTS

### The D253R *syntaxin* mutation affects sleep

Compelling evidence suggests that wakefulness leads to a net synaptic potentiation,^3,5^ and this increase is associated with greater pressure to sleep in order to renormalize (i.e., downregulate) synaptic weight in both humans and animals,^6–8^ including fruit flies.^9–12^ Based on this evidence, we hypothesized that a broad reduction of the synaptic function due to an alteration in neurotransmitter release would lead to reduced synaptic potentiation and the need for downregulation during sleep. As a consequence, if the homeostatic sleep pressure is associated with the necessity of synaptic renormalization, then a reduction in the time spent asleep should be observed.

To test this idea, we analyzed sleep behavior in transgenic flies expressing a mutated form of the SNARE protein Syntaxin1A (*Syx*^*D253R*^), in which the aspartic acid residue 253 was substituted with arginine (D253R). Previously, we showed that *Syx*^*D253R*^ induces a reduction both in the frequency of miniature end plate potentials and in the amplitude of evoked junction potentials at the level of the neuromuscular junction of *Drosophila* larvae.^2^ The reduction of spontaneous and evoked neurotransmitter release following expression of *Syx*^*D253R*^ was not due to the lack of SNARE complex formation, but instead to the probability of formation of a rosette-like super-complex around the point of contact between vesicle and presynaptic membranes.^2^ Therefore, this amino acid substitution seems to interfere with the capacity to mediate vesicle fusion, likely leading to a dampening in the communication among neurons without completely silencing it. Here, we exploited the UAS-Gal4 binary system to alter the balance of the molecular nanomachine regulating synaptic release in the entire adult *Drosophila* nervous system.^13^ By using the pan-neuronal *elav*-Gal4 driver, we expressed either the mutated *Syx*^*D253R*^ or the wild-type (*Syx*^*WT*^) isoform of Syntaxin1A under the control of the UAS promoter in a wild-type background (**Figure 1A**). We then assessed the sleep pattern of 119 *Syx*^*WT*^ flies, 111 *Syx*^*D253R*^ flies and 111 controls (parental flies bearing only the *elav-*Gal4), maintained for 5 days (the first and last day were discarded) in 12h:12h light:dark cycles (**Figure 1B**, left). A video-tracking analysis was additionally performed to evaluate the walking locomotion in greater detail (**Figure 1B**, right).

**Figure 1.**
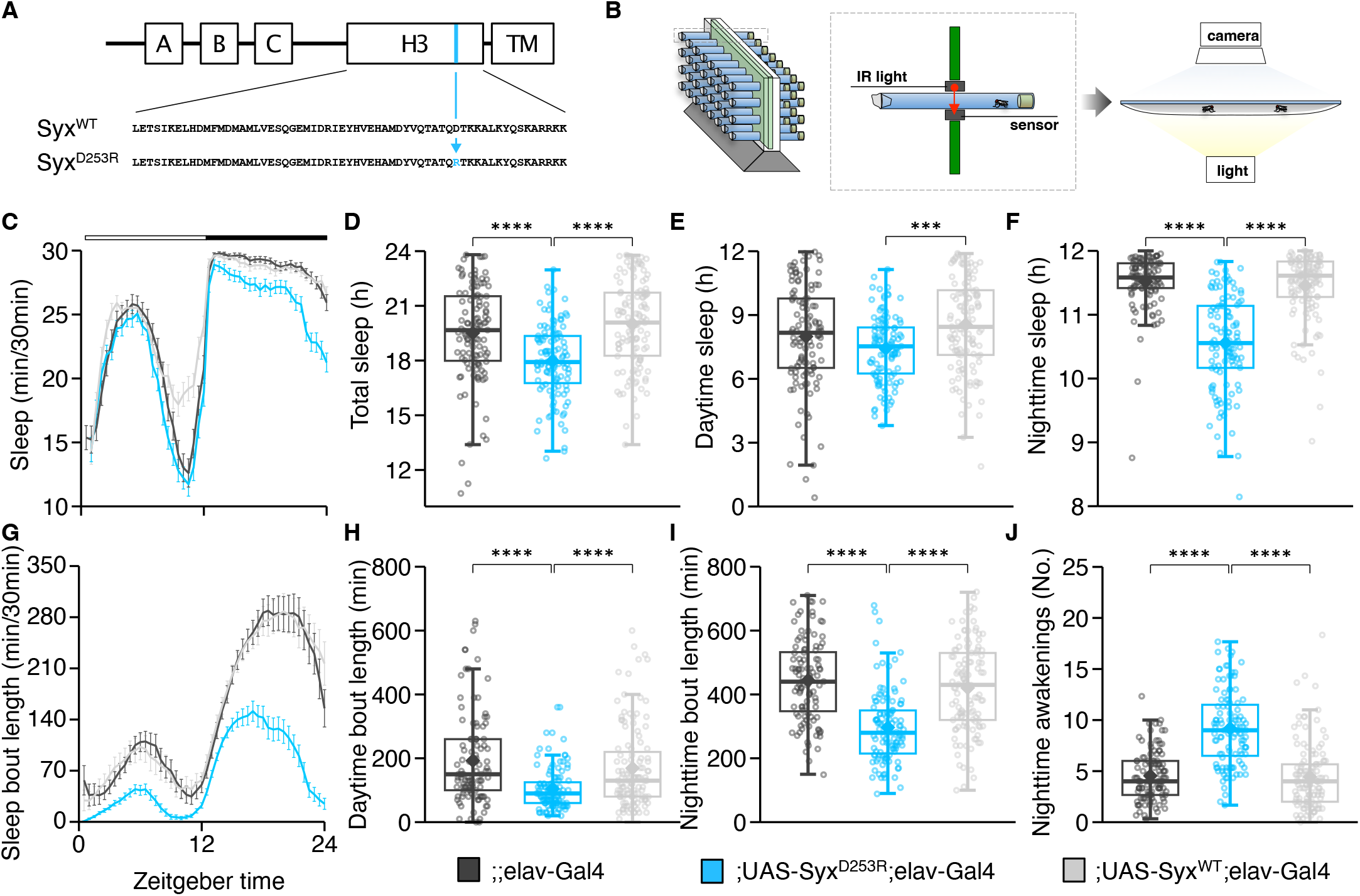
Sleep pattern is compromised in D253R syntaxin mutation. (A) On the left is represented the Syntaxin1A protein with its domains. Under the schematic domains representation, there are the amino acid sequence of the specific H3 domain, an important region for the formation of the SNARE core complex located just before the transmembrane domain (TM): in the Syx wild-type gene and in the mutated form. A, B and C are N-terminal interaction domains. The mutated Syx^D253R^ form has the aspartic acid 253 (D, aspartic acid) substituted with an arginine (R, arginine). On the right is presented the DAM monitor system used to record, through a sensor, the number of times the flies break an infrared (IR) beam positioned at the middle of a glass tube within which the flies were individually loaded. (B) To accurately investigate the locomotor activity of flies, video-tracking recordings were also performed. Flies were loaded in groups of ten within an open-field arena where they were free to walk but not to fly. (C) Average sleep profile in the three genotypes (mean ± SEM). Syx^D253R^ mutant flies slept less than both Syx^WT^ and elav-Gal4 control flies. Repeated-measures ANOVA (rANOVA with Greenhouse-Geisser correction for all analyses) detected a significant interaction effect between time and genotype (F_(14, 2388)_ = 9.17, η^2^=.04, p <.0001). (D) The same rANOVA also detected a main effect for genotype (F_(2, 338)_ = 22.49, η^2^=.04, p <.0001). Syx^D253R^ flies exhibited reduced total sleep time in comparison to Syx^WT^ and elav-Gal4 (Syx^WT^ – Syx^D253R^, p <.0001; Syx^WT^ – elav-Gal4, p =.2527; Syx^D253R^ – elav-Gal4, p <.0001). All analyses were a three-day average. Pairwise post-hoc comparisons were adjusted using Bonferroni method (*p <.05; **p <.01; ***p <.001; ****p <.0001; n = 111-119 flies per genotype). Data are presented as box plots with the box defining first and third quartiles, solid line crossing the box representing the median, and whiskers defining the lowest and the highest values within the 1.5 interquartile range. The rhombus represents the mean and the circles represent the three-day mean values for each fly. (E) Separate one-way ANOVA focused on daytime sleep period detected a main effect for genotype (F_(2, 338)_ = 8.21, η^2^=.05, p =.0003). Syx^D253R^ flies showed reduced sleep duration compared to Syx^WT^ (Syx^D253R^ – elav-Gal4, p =.1171; Syx^WT^ – elav-Gal4, p =.1066; Syx^WT^ – Syx^D253R^, p =.0002). Paired t-test with Bonferroni method were used as in (D). Data are presented as in (D). (F) One-way ANOVA as in (D) for nighttime sleep period detected a genotype effect again (F_(2, 338)_ = 98.27, η^2^=.37, p <.0001). Syx^D253R^ flies exhibited reduced nighttime sleep compared to Syx^WT^ and elav-Gal4 (Syx^D253R^ – elav-Gal4, p < 0001; Syx^WT^ – elav-Gal4, p =.6555; Syx^WT^ – Syx^D253R^, p <.0001). Paired t-test with Bonferroni method were used as in (D). Data are presented as in (D). (G) The graph represents the mean sleep bout duration profile in bins of 30 min (mean ± SEM). A rANOVA detected a significant main genotype effect (F_(2, 338)_ = 36.60, η^2^=.10, p <.0001) and an interaction between time and genotype (F _(8, 1433)_ = 17.13, η^2^=.05, p <.0001). Syx^D253R^ flies showed a general lower sleep bout duration profile than Syx^WT^ and elav-Gal4 flies (Syx^D253R^ – elav-Gal4, p <.0001; Syx^WT^ – elav-Gal4, p =.9853; Syx^WT^ – Syx^D253R^, p <.0001). Post-hoc comparisons as in (D). (H) One-way ANOVA on the maximum daytime sleep bouts duration showed a significant effect for genotype (F_(2, 338)_ = 15.77, η^2^=.09, p <.0001). Syx^D253R^ flies exhibited a reduced sleep bout duration compared to the other two genotypes (Syx^D253R^ – elav-Gal4, p<.0001; Syx^WT^ – elav-Gal4, p =.1858; Syx^WT^ – Syx^D253R^, p =.0005). Post-hoc comparisons as in (D). Data are presented as in (D). (I) As in (G), one-way ANOVA on the maximum nighttime sleep bouts duration showed a significant main genotype effect (F_(2, 338)_ = 47.95, η^2^=.22, p <.0001). Syx^D253R^ flies showed reduced sleep bout duration (Syx^D253R^ – elav-Gal4, p <.0001; Syx^WT^ – elav-Gal4, p =.9736; Syx^WT^ – Syx^D253R^, p <.0001). Data are presented as in (D). (J) One-way ANOVA on nighttime awakenings showed a significant effect of genotype (F_(2, 338)_ =s 82.65, η^2^ =.33, p <.0001). Syx ^D253R^ flies showed increased number of nighttime awakenings compared to the other two genotypes (Syx^D253R^ – elav-Gal4, p <.0001; Syx^WT^ – elav-Gal4, p =.9648; Syx^WT^ – Syx^D253R^, p <.0001). Post-hoc as in (D). Data are presented as in (D). As reported in the legend at the bottom of panel: Syx^D253R^ flies are depicted in light blue, Syx^WT^ in light grey and elav-Gal4 in deep grey.

In line with our hypothesis, the results showed a reduction in the total amount of sleep in *Syx*^*D253R*^ expressing flies compared to both control and *Syx*^*WT*^ expressing flies (**Figure 1C, 1D, 1E** and **1F**). Furthermore, *Syx*^*D253R*^ flies revealed a clear reduction in the average sleep bout duration compared to both *Syx*^*WT*^ and *elav*-Gal4 flies (**Figure 1G**). Finally, *Syx*^*D253R*^ flies showed an average reduction in the maximum sleep bout duration both during daytime and nighttime periods, which is associated with a greater number of nighttime awakenings (**Figure 1H, 1I** and **1J**). These results suggest an impairment of sleep continuity in *Syx*^*D253R*^ flies.

In a previous study, we identified a highly conserved residue (R206) in SNAP-25 using radial modelling of the neuroexocytosis nanomachine which plays a major role in the protein-protein interactions between SNARE complexes for the formation of the super-complex.^2^ Thus, we also tested whether flies pan-neuronally expressing a mutated SNAP-25, bearing an arginine to aspartic acid (R206D) substitution, would show an effect on sleep similar to what was observed by *Syx*^*D253R*^ expressing flies. Surprisingly, we did not detect any significant sleep pattern alterations in these flies compared to controls (**Figure S1A**).

### Locomotor activity is specifically altered during nighttime

To exclude that the sleep defects observed in *Syx*^*D253R*^ were not simply a consequence of a generalized increase in locomotion, we specifically looked at the activity profiles. *Syx*^*D253R*^ individuals showed at a glance higher locomotor activity across the day with respect to controls (**Figure 2A** and **S2A**). However, taking into account only the periods of activity, we did not find significant differences among genotypes in the number of times flies crossed the housing tube per minute, meaning that *Syx*^*D253R*^ flies behave like the controls when active (**Figure 2B**). By analyzing the number of tube crossings every 30 min during the daytime, we found a significant difference only between *Syx*^*D253R*^ and *Syx*^*WT*^ flies (**Figure 2C**). On the other hand, *Syx*^*D253R*^ flies showed a significantly higher level of activity than controls and *Syx*^*WT*^ flies during the nighttime (**Figure 2D**). These data show that *Syx*^*D253R*^ flies are not hyperactive in general and depict a specific effect of the mutation on sleep.

**Figure 2.**
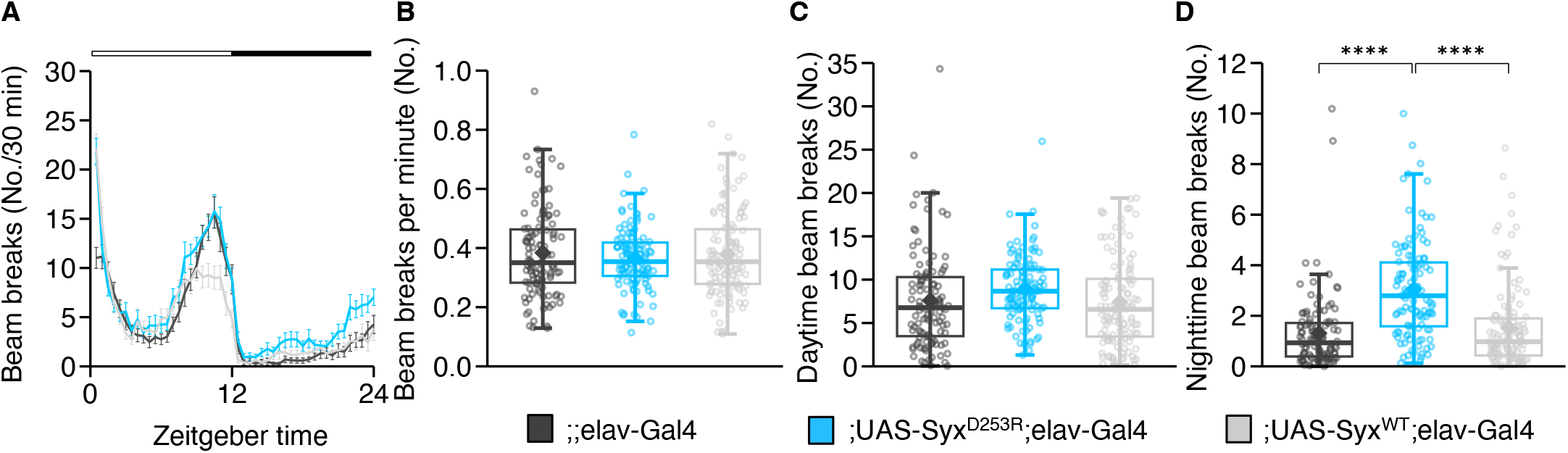
Nighttime locomotor activity is increased in D253R syntaxin mutation. (A) Activity profile measured as beam breaks every 30 minutes (mean ± SEM). Syx^D253R^ flies showed a significant increase in locomotor activity in comparison to the other two genotypes. A rANOVA detected a significant main genotype effect (F_(2, 338)_ = 11.11, η^2^=.01, p <.0001) and an interaction effect between time and genotype (F_(25, 4243)_ = 7.16, η^2^=.03, p <.0001). Pairwise post-hoc revealed a significant difference between Syx^D253R^ and the other two genotypes (Syx^WT^ – Syx^D253R^, p =.0001; Syx^WT^ – elav-Gal4, p =.9995; Syx^D253R^ – elav-Gal4, p =.0002). (B) Syx^D253R^ flies exhibited no evidence of hyperactivity, as measured by beam breaks counts during a daytime minute of activity, relative to Syx^WT^ and elav-Gal4 controls. One-way ANOVA did not detect a significant genotype effect (F_(2, 338)_ = 0.35, η^2^ =.002, p =.71). Data are presented as box plots with the box defining first and third quartiles, solid line crossing the box representing the median, and whiskers defining the lowest and the highest values within the 1.5 interquartile range. The rhombus represents the mean and the circles represent the three-day mean values for each fly. (C) During the daytime period Syx^D253R^ flies exhibited more activity than the Syx^WT^ flies. One-way ANOVA detected a significant genotype effect (F_(2, 338)_ = 3.26, η^2^=.02, p =.04) and the pairwise post-hoc comparisons showed significant difference only between Syx^D253R^ and Syx^WT^ flies (Syx^WT^ – Syx^D253R^, p =.0487; Syx^WT^ – elav-Gal4, p =.9535; Syx^D253R^ – elav-Gal4, p =.1053). Data are presented as in (B). (D) During nighttime Syx^D253R^ flies exhibited more activity than the Syx^WT^ and elav-Gal4 controls (Syx^WT^ – Syx^D253R^, p <.0001; Syx^WT^ – elav-Gal4, p =.6054; Syx^D253R^ – elav-Gal4, p <.0001). One-way ANOVA showed a significant genotype effect (F_(2, 338)_ = 35.01, η^2^=.17, p <.0001). Data are presented as in (B). All analyses were a three-day average and pairwise post-hoc comparisons were adjusted with Bonferroni method (*p <.05; **p <.01; ***p <.001; ****p <.0001; n = 111-119 flies per genotype). As reported in the legend at the bottom: Syx^D253R^ flies are depicted in light blue, Syx^WT^ in light grey and elav-Gal4 in deep grey.

### A finer kinematics analysis confirms normal walking in the daytime

Given that the system used for the analysis of sleep/wake patterns tends to underestimate the amount of movement occurring during bouts of activity,^14,15^ we decided to address this issue specifically by using a video-tracking setup. We thus recorded the activity of 59 *Syx*^*WT*^, 51 *Syx*^*D253R*^ and 35 control male flies in an open-field arena for a relatively short period of time (i.e., 10 min). Inside the arena, designed to confine flies in a 2D space, individuals were free to walk and interact with each other, but could not fly. After video recording the flies’ behavior, we tracked and analyzed offline several parameters associated with the flies’ walking kinematics. The total number of stops and walking bouts made by *Syx*^*D253R*^ flies did not show significant differences compared to *Syx*^*WT*^ and *elav*-Gal4 flies (**Figure 3A** and **3B**). The same was observed for the length of walking bouts (**Figure 3C**), total path length travelled (**Figure 3D**), number of sharp turns (**Figure 3F**, see Methods), and forward velocity (**Figure 3D**). Overall, *Syx*^*D253R*^ flies did not show abnormal walking locomotion compared to controls.

**Figure 3.**
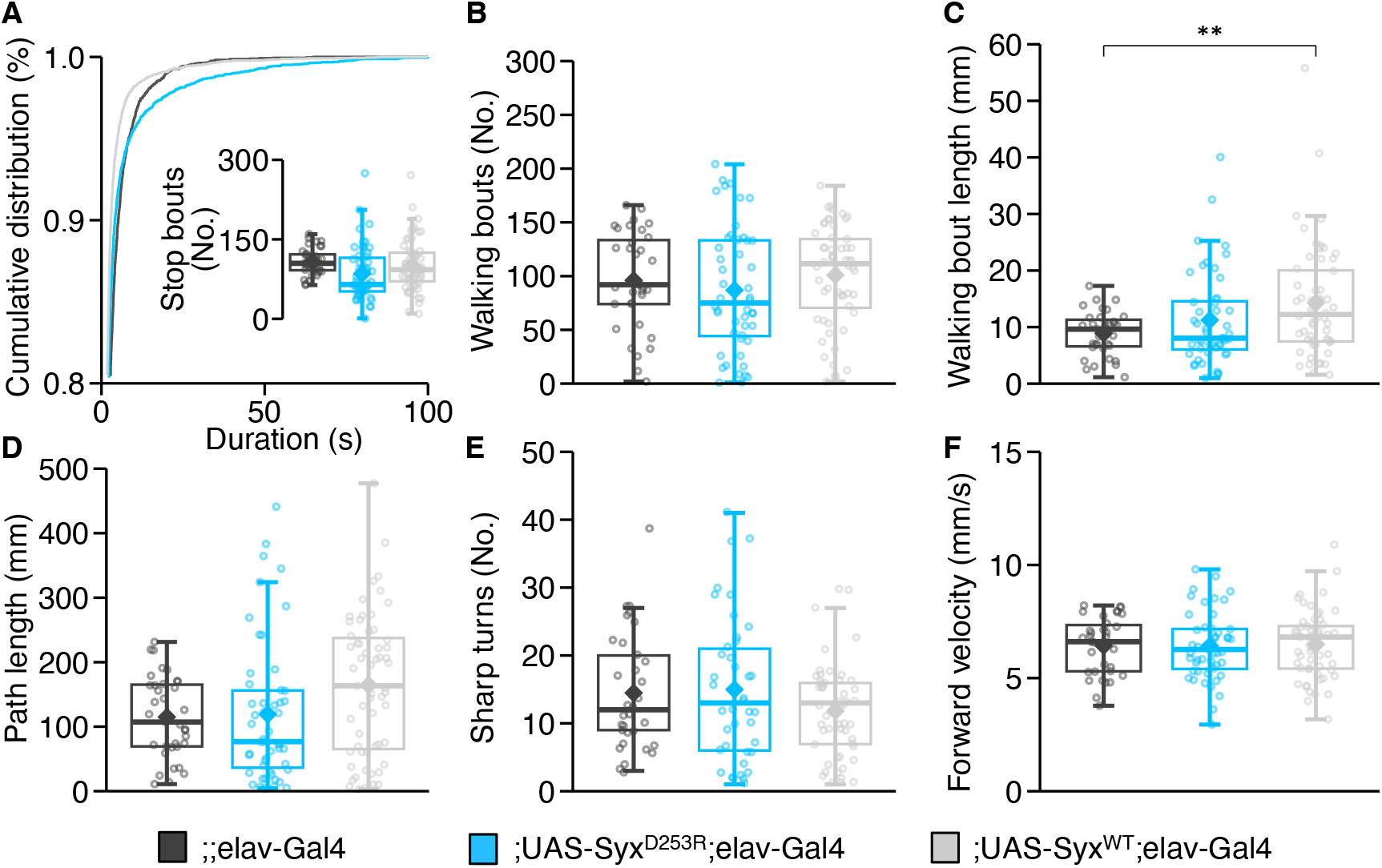
Normal daytime walking behavior in D253R syntaxin mutation. (A) Cumulative distribution of the stop bouts duration per genotype. On the bottom right, the average number of stops is represented. One-way ANOVA did not detect a significant genotype effect (F_(2, 142)_ = 2.95, η^2^ =.04, p =.06). Data are presented as box plots with the box defining first and third quartiles, solid line crossing the box representing the median, and whiskers defining the lowest and the highest values within the 1.5 interquartile range. The rhombus represents the mean and the circles represent the values for each fly. (B) Average number of walking bouts in the three genotypes, as highlighted in the methods details. One-way ANOVA did not detect a significant genotype effect (F_(2, 136)_ = 0.99, η^2^=.01, p =.37). Data are presented as in (A). (C) Average walking bout length in the three genotypes. One-way ANOVA showed a significant genotype effect (F _(2, 136)_= 4.77, η^2^=.07, p =.01) and the pairwise post-hoc comparisons showed significant difference only between elav-Gal4 and Syx^WT^ (Syx^WT^ – Syx ^D253R^, p =.1532; Syx^WT^ – elav-Gal4, p =.0085; Syx^D253R^ – elav-Gal4, p =.4107). Data are presented as in (B). (D) Box plot per genotype of path length travelled by flies in ten minutes of video-tracking recording. One-way ANOVA detected a significant genotype effect (F _(2, 142)_ = 3.83, η^2^=.05, p =.02) but pairwise post-hoc comparisons did not show any significant difference among genotypes (Syx^WT^ – Syx^D253R^, p =.0520; Syx^WT^ – elav-Gal4, p =.0587; Syx^D253R^ – elav-Gal4, p =.9806). Data are presented as in (B). (E) Average number of sharp turns (as described in methods) in the three genotypes. One-way ANOVA did not detect a significant genotype effect (F_(2, 123)_ = 1.78, η^2^=.03, p =.01). Data are presented as in (B). (F) Box plot of the average forward velocity during recording. One-way ANOVA did not detect a significant genotype effect (F_(2, 136)_ = 0.04, η^2^ =.0006, p =.96). Data are presented as in (B). In all analyses, pairwise post-hoc comparisons were adjusted with Bonferroni method (*p <.05; **p <.01; ***p <.001; ****p <.0001; n = 51 Syx^D253R^; n = 59 Syx^WT^; n = 35 elav-Gal4). As reported in the legend at the bottom: Syx^D253R^ flies are depicted in light blue, Syx^WT^ in light grey and elav-Gal4 in deep grey.

### The circadian rhythm is maintained

The evidence of sleep alterations in *Syx*^*D253R*^ mutants does not rule out possible changes in their circadian rhythm. To address this point, we recorded the locomotor activity of flies in constant darkness (12h:12h dark:dark) following a three-day entrainment period, and analyzed their free-running locomotor activity. Although the overall activity was severely reduced with respect to the entrained 12h:12h light:dark condition (**Figure 4A** and **4B**), the locomotor circadian rhythm was maintained in all genotypes (**Figure S3**). The circadian period, as well as the phase, of *Syx*^*D253R*^ flies did not differ significantly from controls (**Figure 4C**). The lack of significant changes in the circadian rhythm of *Syx*^*D253R*^ flies suggests that the mutation does not impinge on the circadian clock,^16^ but rather on the homeostatic sleep-driving process.^17^

**Figure 4.**
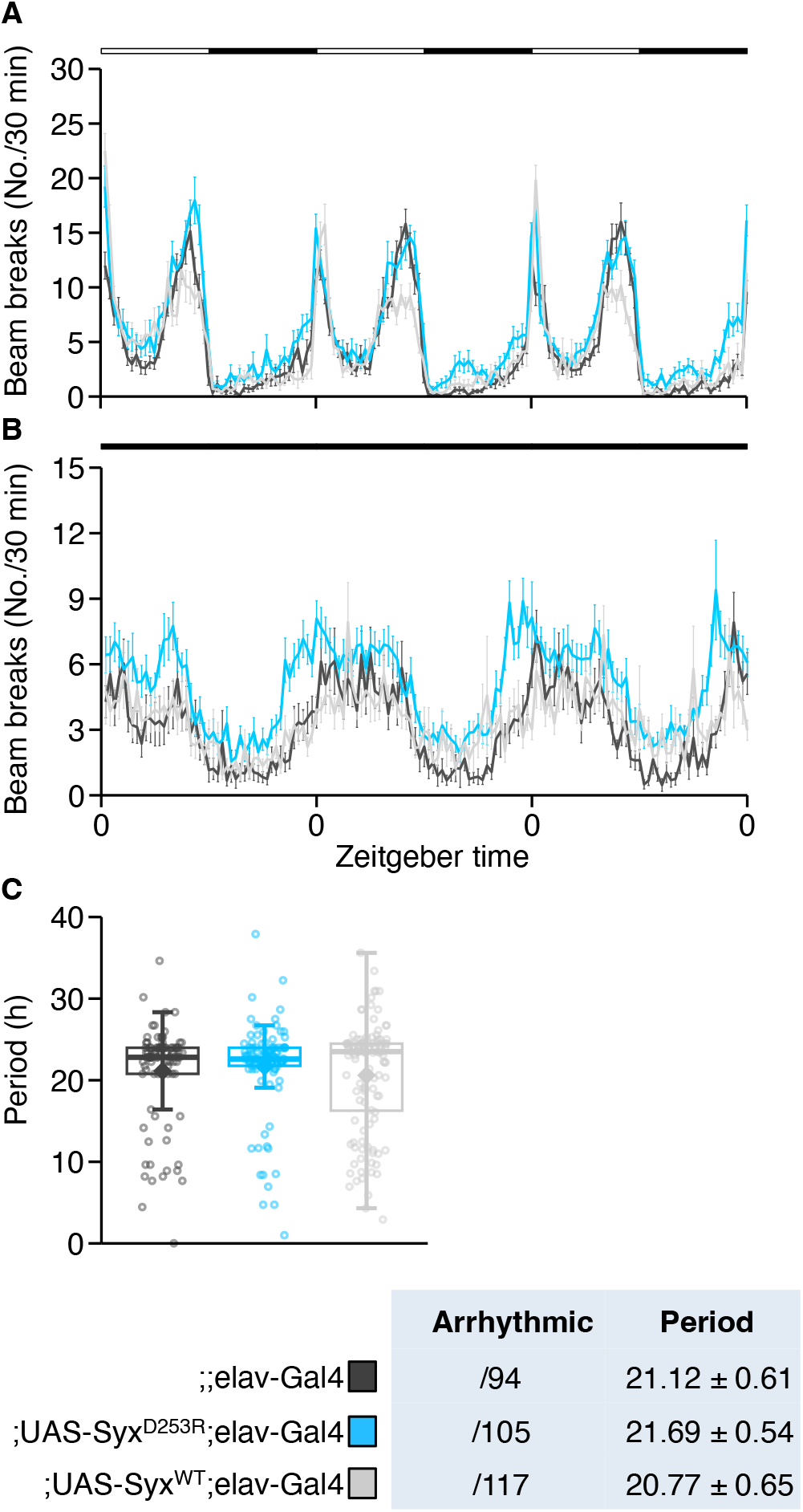
Unaltered circadian rhythmicity in D253R syntaxin mutation. (A) The graph represent the average number of beam breaks every 30 minutes during 3 days of 12h:12h light:dark (LD) condition (mean ± SEM). (B) Circadian locomotor activity (i.e., average number of beam breaks every 30 minutes) during 3 days of 12h:12h dark:dark condition. (C) In the table are reported the number of arrhythmic flies for genotype and the mean of circadian period (mean ± SEM). Syx^D253R^ flies did not exhibit change in circadian rhythmicity compared to the other two genotypes during 3 days of 12h:12h dark:dark condition. One-way ANOVA did not detect a significant genotype effect on circadian period (F_(2, 313)_ = 0.62, η^2^ =.004, p =.54). Data are presented as box plots with the box defining first and third quartiles, solid line crossing the box representing the median, and whiskers defining the lowest and the highest values within the 1.5 interquartile range. The rhombus represents the mean and the circles represent the three-day mean values for each fly. As reported in (C), in all images: Syx^D253R^ flies are depicted in light blue, Syx^WT^ in light grey and elav-Gal4 in deep grey.

## DISCUSSION

Working under the synaptic homeostasis hypothesis,^3^ a previous study has demonstrated the role of the ELKS-family active zone core scaffold protein, Bruchpilot, which seems to be driving part of the active zone plasticity, in promoting sleep rebound.^11^ In this study, we focused on an alteration in the synaptic exocytosis nanomachine selectively impacting sleep. Specifically, an amino acid replacement in Syntaxin1A induced fragmented and reduced amounts of night sleep. Syntaxin1A is part of the tripartite SNARE complex whose integrity is key for neuroexocytosis. Clear evidence of its importance is the terrible effects of botulinum (BoNT) and tetanus toxins on the nervous system, which completely block neurotransmitter release by cleaving one or more components of the SNARE complex. The SNARE complexes bind together around the point of contact between the vesicle and neuronal membrane forming a rosette-like super-complex.^18–20^ If the formation of this super-complex is prevented, the synaptic exocytosis nanomachine might not be able to clear the energy step necessary for membrane fusion and neurotransmitter release. This could explain why the isotype A of BoNT (BoNT/A) retains its physiological consequences despite the cleaving of the last residues of SNAP-25 C-terminal region that do not interfere with the assembly of the SNARE complex. According to model predictions,^2,18,21^ one of the SNAP-25 C-terminal residues cleaved by BoNT/A (i.e., R206 in *Drosophila*) plays a key role in forming a protein-protein interaction with the Syntaxin1A (i.e., residue D253) of a neighboring SNARE complex. This interaction would allow the SNARE complexes to connect with each other in a radial arrangement. Electrophysiological experiments in *Drosophila* larval neuromuscular junction have confirmed these residue substitutions hinder the super-complex formation and result in a reduction of both evoked and spontaneous synaptic release.^2^ A small group of cytoplasmic proteins called complexin and some general anesthetics might also act on the formation of the super-complex.^18,22,23^

*Syx*^*D253R*^ mutated protein expression seems to strongly compromise sleep given that wake activity is not accompanied by changes in locomotion. These sleep alterations do not appear to be associated with impairments of the circadian rhythm. How does the *Syx*^*D253R*^ mutation affect homeostatic sleep? The average reduction of evoked junction potentials in *Syx*^*D253R*^ transgenic flies has been estimated to be around 40%, meaning that this genetic manipulation does not completely block the synaptic activity.^2^ This explains why our flies did not show a more severe behavioral phenotype such as muscle paralysis, as indeed observed in all BoNT isotypes envenomation, except BoNT/A.^24^ Though, the widespread distribution of a diminished synaptic function could be sufficient to ease connectivity and metabolic load for neuronal processing, especially during daytime. Therefore, in line with the synaptic homeostasis hypothesis,^3^ the amount of sleep necessary to restore cellular functions and renormalize synaptic strengths across the brain could be reduced accordingly. Noteworthy is the increase by 30% in the expression level of Syntaxin1A after 24h of sleep deprivation.^10^

Interestingly, the substitution of threonine 254 with isoleucine in Synatixn1A (*Syx*^*T254I*^) provokes an increase of both spontaneous and evoked neurotransmitter release at the neuromuscular junction of third instar larvae.^25,26^ Similarly, ON/OFF transients in electroretinograms and action potentials (evoked by electrical stimulation of the giant fiber) in the indirect flight muscles seem to be preserved or increased in adult flies that carry the *Syx*^*T254I*^ mutation.^26^ These mutant flies show a severe paralytic behavior which is thought to be the consequence of an exacerbated vesicle depletion caused by an increased synaptic activity.^24,26^ A similar behavior has been observed in CAKI mutant flies where the spontaneous vesicle release is increased.^27^ Unfortunately, there are no data available on sleep in *Syx*^*T254I*^ mutants, probably because heterozygous flies still show a paralytic phenotype. However, flies expressing the *Syx*^*T254I*^ transgene pan-neuronally (using *elav*-Gal4 as driver) have shown an increased sensitivity to isoflurane anesthesia.^28^ Yet, a negative correlation between isoflurane resistance and sleep has been suggested, implying that the reduced isoflurane resistance of *Syx*^*T254I*^ expressing flies might be accompanied by an increased amount of sleep.^28^ Taken together, these results hint at a role of Syntaxin1A in regulating the need for sleep.

Adult flies expressing a mutated form of SNAP-25 (*SNAP*^*R206D*^), where arginine 206 is substituted with aspartic acid (R206D), did not show alterations in sleep. This is despite the fact that, as mentioned above, these transgenic flies exhibit both a decrease in miniature and evoked post-synaptic potentials as observed in *Syx*^*D253R*^ transgenic flies.^2,21^ Unlike Syntaxin1A, SNAP-25 is thought to affect synaptic transmission not only pre-synaptically, as a critical component of the SNARE complex, but also post-synaptically.^29,30^ Through an interaction mediated by the C-terminal nine amino acids, SNAP-25 seems to favor the activity-stimulated endocytosis of post-synaptic glutamate receptors and the subsequent triggering of a form of long-term depression.^29,30^ This opposite pre– and post-synaptic role of SNAP-25 could explain the seemingly contradictory result on sleep. Namely, the SNAP-25 mutants studied here could compensate for a faulty vesicle release machinery with a less sensitive negative regulatory mechanism. The net effect would translate to “normal” synaptic weight requiring “normal” downregulation during sleep.

Sleep alterations, defined as modifications in total amount of sleep, sleep fragmentation, or daytime sleep are commonly found in people with psychiatric disorders.^31^ Alterations of the SNARE complex may lead to synaptic dysfunctions that undermine sleep behavior and cognitive activity.^32^ Several studies support the idea that the expression of Syntaxin1A could be altered in ADHD, autism spectrum disorders and schizophrenia.^33–36^ Data from vertebrate and invertebrate model organisms seem to corroborate a correlation between Syntaxin1A and behavioral alterations. In mice, 6

Syntaxin1A knockout has been related to behavioral abnormalities,^37^ while in bees, Syntaxin1A polymorphisms have been associated with alteration of social behavior.^38^ Here, we studied the impact on sleep of a pan-neuronal alteration of Syntaxin1A in flies, confirming its putative role in the regulation of sleep homeostasis. However, other complex behaviors, such as social interaction, or microbehaviors that we were unable to resolve could be affected by this alteration. Driving panneuronal modifications of key synaptic proteins in fruit flies, combined with high-resolution video analyses,^39^ will be extremely informative to clarify how these proteins influence specific behaviors.

### Contact for Reagent and Resource Sharing

Further information and requests for resources and reagents should be directed to and will be fulfilled by the Lead Contact, Aram Megighian.

### Experimental Model and Subject Details

#### Fly Strains

We used the driver line C155, *elav*-Gal4, from Bloomington Drosophila Stock Center (Bloomington, IN). The UAS-*Syx*^*D253R*^ and UAS-*Syx*^*WT*^ lines were made by our laboratory as previously described.^2^ Flies were raised in standard yeast-glucose-agar medium and maintained at 25°C, 70% relative humidity, and in a 12-hour light:12-hour dark cycle. Experiments were carried out with adult male flies (3-5 days old).

### Method Details

#### Sleep Assay

Fly locomotor activity was monitored continuously for three days under 12h:12h light:dark condition at 25°C by using the Drosophila Activity Monitoring System 2 (DAM2, Trikinetics, Waltham, MA). Flies were left to acclimatize for a complete day before starting the locomotor activity recording. The output from the DAM2 consists of the number of times a fly crosses a set of infrared beams in a period of 1 min (bin). A sleep episode (bout) is defined as 5 or more consecutive bins of immobility.^40^ These data were subsequently processed and analyzed with custom scripts by using the integrated development environment RStudio (version 1.1.463),^41^ running on the R statistical computing software (version 3.5.1).^42^

#### Circadian Analysis

As for the sleep analysis, male flies housed in a 65 mm glass tube were recorded for three continuous days under 12h:12h dark:dark condition by using the DAM2 system. Flies were entrained for three days in 12h:12h light:dark cycles before switching to constant darkness. The data in output from the DAM2 were subsequently analyzed with the ActogramJ plugin^43^ for ImageJ (National Institutes of Health).^44^ The Lomb-Scargle periodogram was conducted for each fly and the data exported for statistical analysis (i.e., one-way ANOVA) performed in RStudio.^41^

#### Locomotion in open-field arena

Walking fly locomotion was investigated in an open-field arena of 10 cm diameter custom-designed to confine flies in 2D space, while impeding flight through the presence of a glass “ceiling”. In each experiment, ten male flies of the same genotype were inserted into the arena. The acrylic glass arena was in turn placed on a platform made of the same material which was located inside a box and lit from below with cool white (6500K) LEDs (Elcart, Milan, IT). A fan maintained a temperature-controlled environment by blowing room air under the platform. A layer of opaque paper placed between the arena and the platform guaranteed a diffuse and homogeneous illumination. The background surrounding the arena contained no visual landmarks. After the flies were loaded into the arena by using a mouth aspirator, they were left to acclimatize in complete darkness for 5 min before turning on the light. Simultaneously with lights-on, the recording was triggered and the video acquired using a CCD camera (Chameleon 2, Point Grey Research Inc, Richmond, BC, Canada) fitted with a 2.7-13 mm varifocal lens (Fujifilm, Tokyo, JP) that streamed to a desktop computer via USB. Video recordings were acquired at 15 frames/s with a resolution of 1296 x 964 pixels. Each recording lasted 10 min and was acquired by using FlyCapture software (Point Grey Research Inc, Richmond, BC, Canada). All experiments were conducted between zeitgeber time 2 and 4 at a room temperature ranging between 22 and 23°C.

#### Fly tracking and analysis

In order to track the fly positions, the videos recorded were analyzed off-line by using the open-source tracking software CTRAX (California Institute of Technology Fly Tracker).^45^ Next, a series of parameters (e.g., velocity and acceleration) were computed using a MATLAB toolbox provided within CTRAX (i.e., BehavioralMicroarray). These parameters were then imported into RStudio^41^ for further data analysis and statistical computing. We classified the fly locomotor activity in three simple components: bouts of walking, stopping and sharp turning. The walking behavior was defined as a bout of walking with velocity and duration equal to or higher than 2.5 mm/s and 250 ms, respectively. Stopping was described as a bout in which the walking velocity was equal or lower than 1 mm/s and its duration was equal to or longer than 250 ms. Finally, the sharp turning was categorized as a turning with angular velocity no slower than 60 deg/s and duration of at least 320 ms. By using these criteria, we calculated the number of times each fly initiated to walk, the bout length, the number of stops, and the number of sharp turns. The average of forward velocity and acceleration were also computed during walking bouts.

#### Quantification and Statistical Analysis

Repeated-measures and one-way analysis of variance (ANOVA) were performed using the R package *afex*.^46^ Multiple comparisons were performed by using post hoc pairwise t-tests adjusted with the Bonferroni method.

## Data and Software Availability

The datasets and code generated during this study are available upon request.

## Acknowledgements

We thank Dr. Paola Cisotto for her unconditional technical support.

## Author Contributions

Conceptualization, G.F., M.Z. and A.M.; Formal Analysis, G.F.; Investigation, G.F. and A.M.; Resources, A.M.; Writing – Original Draft, G.F. and N.C.; Writing – Review & Editing, G.F., N.C., G.M.M., M.Z. and A.M.; Supervision, M.Z. and A.M; Funding Acquisition, M.Z. and A.M.

## Supplemental Figures

**Figure S1.**
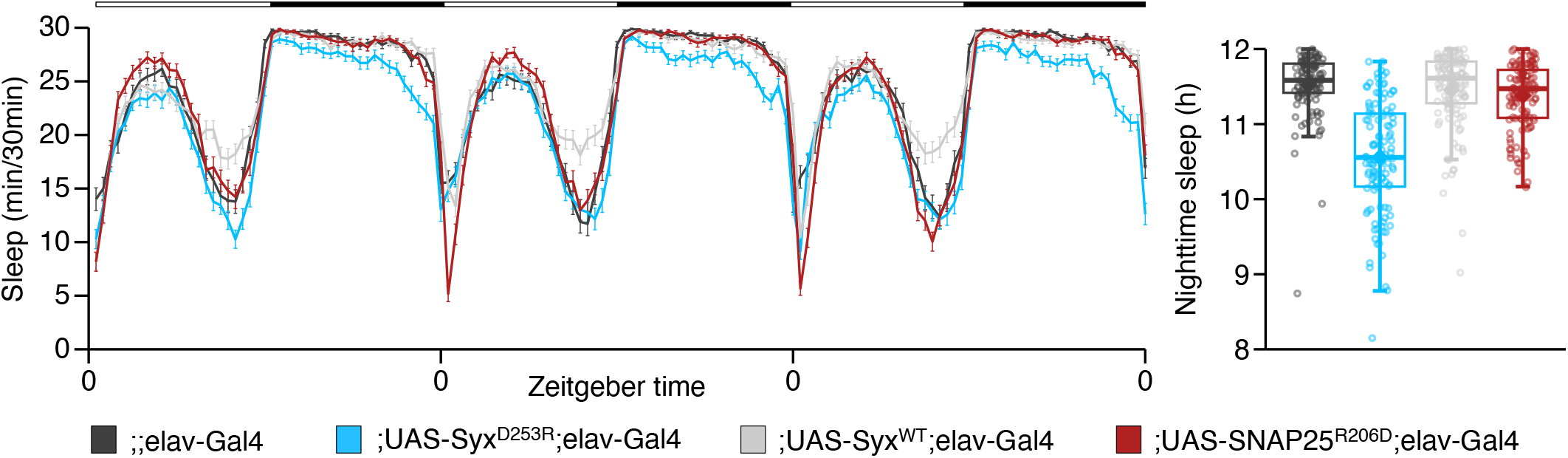
Sleep is not compromised in SNAP^R206D^. On the left, sleep profile in SNAP^R206D^, Syx^D253R^, Syx^WT^ and elav-Gal4 male flies throughout 3 days (mean ± SEM). As reported in the legend at the bottom of image: SNAP^R206D^ flies is depicted in red, Syx^D253R^ in light blue, Syx^WT^ in light grey and elav-Gal4 in deep grey. On the right, box plot of the nighttime sleep in the same genotypes. The box defines first and third quartiles, solid line crossing the box represents the median, and whiskers defines the lowest and the highest values within the 1.5 interquartile range. The rhombus represents the mean and the circles represent the three-day mean values for each fly. One-way ANOVA detected a main effect for genotype (F_(3, 454)_ = 77.83, η^2^=.34, p <.001). SNAP^R206D^ flies exhibited normal amount of total sleep compared to Syx^WT^ and elav-Gal4 (Syx^WT^ – SNAP^R206D^, p = 1; SNAP^R206D^ – elav-Gal4, p =.2867). This analysis was a three-day average and pairwise post hoc comparisons adjusted with Bonferroni’s correction were used (n = 111-119 flies per genotype).

**Figure S2.**
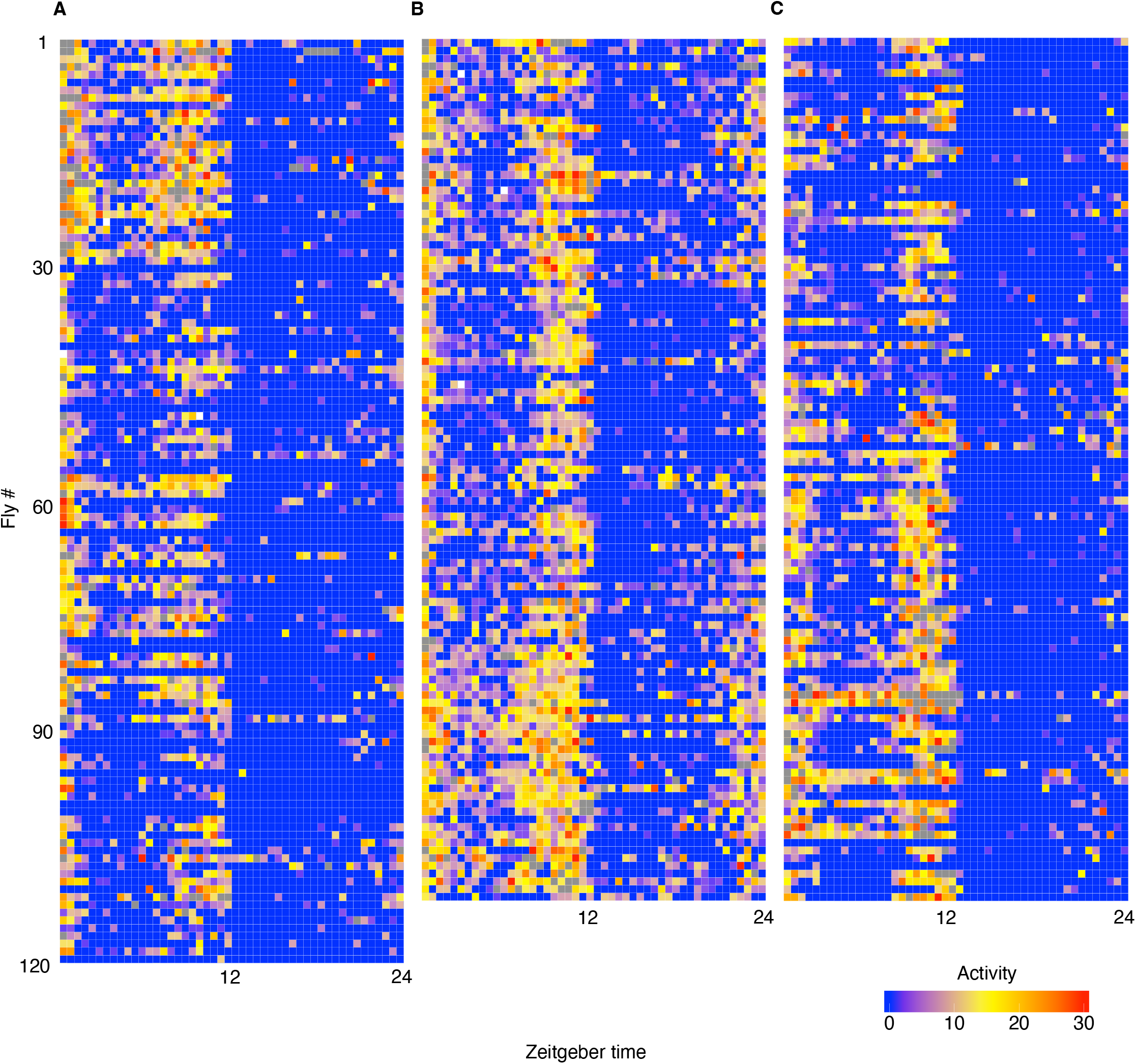
Heatmap of a day average activity in 12h:12h L:D. The figure showed a day average activity in bins of 30 min (in columns) of every fly (in row), in the three tested groups (A, B, C blocks). The colors represent the activity (i.e., the number of beam breaks) as reported in the bottom right legend. (A) Average activity in Syx^WT^ flies every 30 minute (n= 119 flies). (B) Average activity in Syx^D253R^ flies (n= 111). (C) Average activity in elav-Gal4 flies (n= 111).

**Figure S3.**
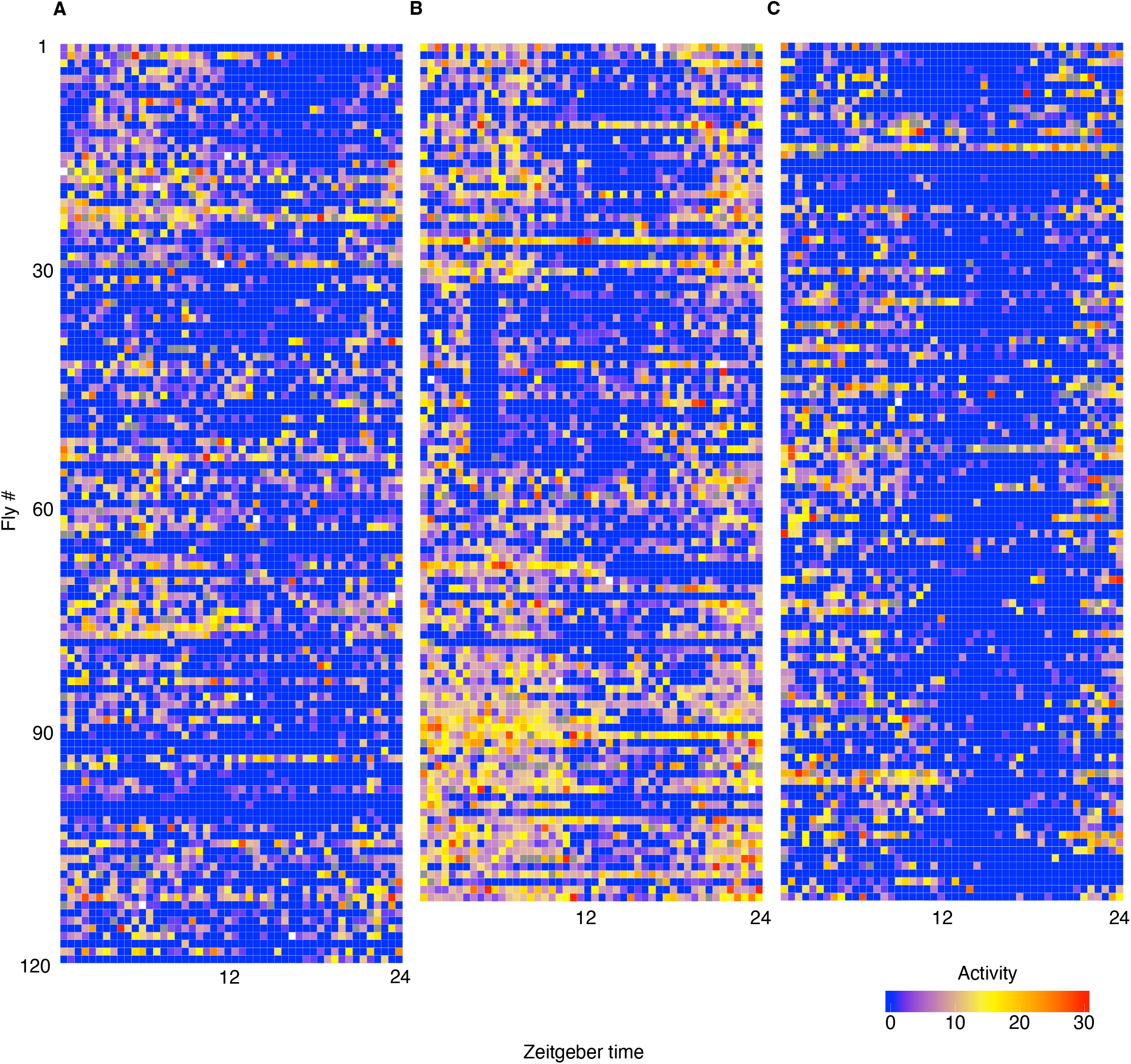
Heatmap of a day average activity in 12h:12h D:D. The figure showed a day average activity in bins of 30 min (in columns) of every fly (in row), in the three tested groups (A, B, C blocks). The colors represent the activity (i.e., the number of beam breaks) as reported in the bottom right legend. (A) Average activity in Syx^WT^ flies every 30 minute (n= 119 flies). (B) Average activity in Syx^D253R^ flies (n= 111). (C) Average activity in elav-Gal4 flies (n= 111).

